# Task-related hemodynamic responses in human early visual cortex are modulated by task difficulty and behavioral performance

**DOI:** 10.1101/2021.07.14.452398

**Authors:** Charlie S. Burlingham, Minyoung Ryoo, Zvi N. Roth, Saghar Mirbagheri, David J. Heeger, Elisha P. Merriam

## Abstract

Early visual cortex exhibits widespread hemodynamic responses in the absence of visual stimulation, which are entrained to the timing of a task and not predicted by local spiking or local field potential (LFP). Such task-related responses (“TRRs”) covary with reward magnitude and physiological signatures of arousal. It is unknown, however, if TRRs change on a trial-to-trial basis according to behavioral performance and task difficulty. If so, this would suggest that TRRs reflect arousal on a trial-to-trial timescale and covary with critical task and behavioral variables. We measured fMRI-BOLD responses in the early visual cortex of human observers performing an orientation discrimination task consisting of separate easy and hard runs of trials. Stimuli were presented in a small portion of one hemifield, but the fMRI response was measured in the ipsilateral hemisphere, far from the stimulus representation and focus of spatial attention. TRRs scaled in amplitude with task difficulty, behavioral accuracy, reaction time, and lapses across trials. These modulations were not explained by the influence of respiration, cardiac activity, or head movement on the fMRI signal. Similar modulations with task difficulty and behavior were observed in pupil size. These results suggest that TRRs reflect arousal and behavior on the timescale of individual trials.

## Introduction

Widespread hemodynamic responses, time-locked to trial onsets, occur in the absence of a visual stimulus in early visual cortex, the earliest site of visual cortical processing (1). These ‘task-related’ responses (hereafter, “TRRs”) have been reported in awake macaques (1–3) using intrinsic-signal optical imaging, and in humans using fMRI (4–6). Unlike stimulus-evoked responses in V1, they are poorly predicted by changes in mean firing rates or LFP amplitudes (1), and unlike spatial attentional responses (6–9), they are spatially diffuse, extending far beyond the focus of spatial attention (1, 6). Previous studies demonstrate that TRRs are modulated in amplitude by reward and correlate with heart rate and pupil size (3, 6), suggesting that they reflect arousal. These findings raise the question of whether TRRs in early visual cortex change from trial to trial with task difficulty and behavioral performance, variables linked to arousal level (10).

In this study, we measured TRRs in human early visual cortex while observers performed a visual orientation discrimination task varying in difficulty, building on the task protocol introduced by Roth et al. 2020 (6). Roth et al (6) averaged fMRI responses over many trials and observed a positive relationship between reward magnitude and trial-averaged TRR amplitude. Here, we update and extend those results by using a general linear mixed model (GLMM) to probe the relation between behavioral performance and response amplitude across trials (11). We found that TRRs in early visual cortex scaled in amplitude with task difficulty, accuracy, reaction time, and lapses, with decreasing strength ascending the visual cortical hierarchy. We found similar modulations in pupil size and cardiac activity, supporting the hypothesis that TRRs are linked to arousal. Our results demonstrate the existence of a widespread hemodynamic response in early visual cortex that reflects arousal on the timescale of individual trials.

## Results

### Task protocol and hypothesis

Human observers (N = 13) performed a visual orientation discrimination task consisting of separate easy and hard runs of trials while undergoing fMRI scanning (Fig 1). Each trial was 15 seconds, composed of a 0.2 s stimulus presentation and 14.8 s inter-stimulus interval. On runs of easy trials, the tilt of the stimulus (grating; diameter, 1.5°) was fixed at ±20° from vertical, yielding discrimination accuracy of ~90%. On runs of hard trials, the tilt was set adaptively with a staircase that yielded ~75% accuracy, typically converging to a tilt of ±1–4° from vertical. The fixation cross was a different color for easy and hard runs (green or red) to ensure that observers were aware of the difficulty of the task. The stimulus was always presented in the same location in the lower right hemifield such that stimulus-evoked responses and spatial attention responses were confined to a small region of the contralateral hemisphere. fMRI responses were measured in the right hemisphere, ipsilateral to the stimulus. These responses in the ipsilateral hemisphere were termed “task-related” because they were neither stimulus-evoked nor related to spatial attention (6). Task-related responses were estimated by averaging over all voxels on the cortical surface in ipsilateral V1. We first used linear regression to project out the global signal from each voxel in V1, a procedure which has been shown to best reduce the influence of physiological processes on the fMRI signal (6, 12). To account for any possible residual physiological artifacts, we additionally predicted cardiac- and respiration-evoked fMRI activity (from cardiac and respiration signals measured during scanning) and projected it out of the fMRI data. We used a GLMM (11) to assess the hypothesis that TRRs scale in amplitude with task difficulty and behavioral performance, while controlling for inter-observer differences in the fMRI signal.

**Fig. 1.**
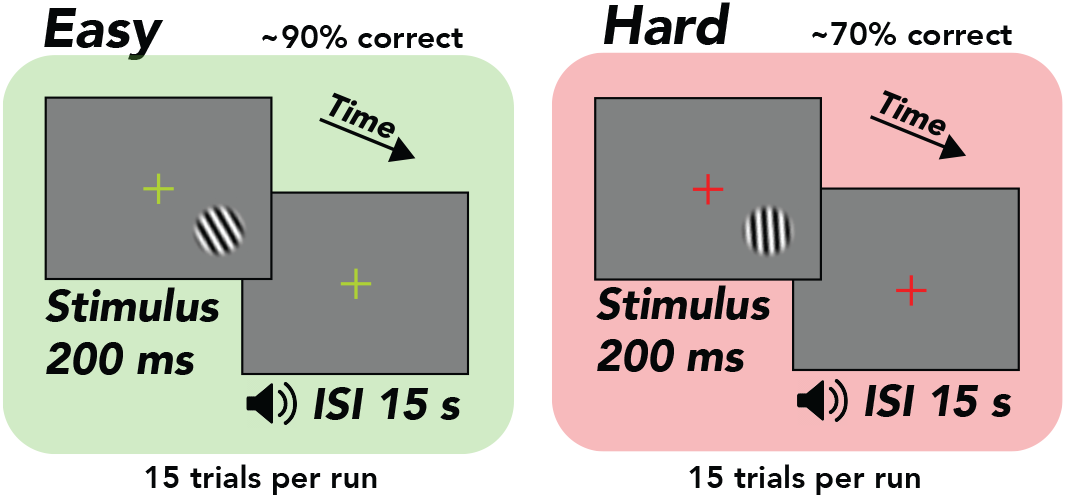
Task, orientation discrimination around vertical. Trial structure, 200 ms stimulus presentation, followed by a 14.8 second inter-stimulus interval, during which the observer made a button press response and immediately received tone feedback. Design, alternation between separate easy (~90% correct) and hard (~70% correct) runs comprising 15 trials each. Stimulus, tilted grating in a raised cosine aperture (diameter of 1.5°, but enlarged for illustrative purposes). Fixation cross, changed colors from green to red, indicating easy and hard runs, respectively.

### Task-related fMRI responses were spatially extensive and entrained to task timing

We observed extensive fMRI responses, extending past the boundaries of atlas-defined V1 (Fig. 2A) and into other visual areas in the occipital, posterior parietal, and temporal cortex. We restricted the analysis to responses in ipsilateral V1, which corresponded to the hemifield in which no stimulus was presented (Fig. 2A). These responses were entrained to trial timing (Fig. 2B), i.e., the maximum amplitude component of the frequency response in ipsilateral V1 was 1/15 Hz, matching the frequency of the task (Fig. 2C). Observers were instructed to covertly attend the peripheral stimulus, which was offset 5 degrees from the central fixation cross. Thus, based on numerous prior studies (6–9), it was assumed that the attention field was restricted to a small region in the contralateral hemisphere surrounding the location of the stimulus. The spatially-extensive activity we observed suggests that TRRs were separate from stimulus-evoked and spatial attention responses.

**Fig. 2.**
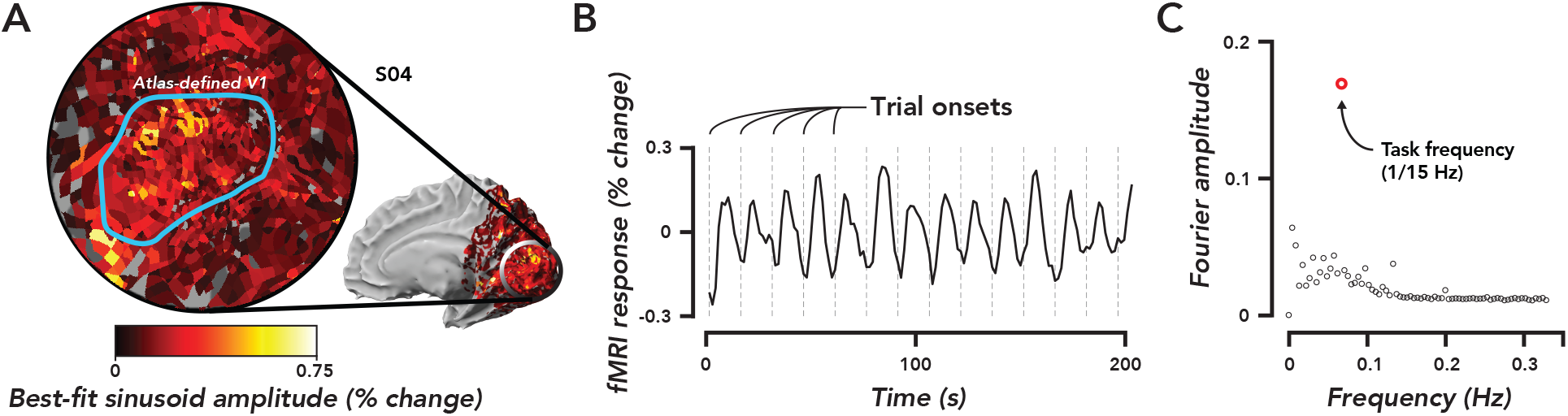
Widespread task-related fMRI responses in ipsilateral V1 were entrained to task timing. **A**. Flat map centered on occipital pole showing fMRI responses. Inset, medial view of right hemisphere, indicated the region of cortex displayed in the flat map. Color map, amplitude of best-fitting sinusoid at the task frequency (1/15 Hz) in units of percent signal change for a single observer (O4). Blue ROI, atlas-defined V1, based on observer’s anatomy. **B**. Time course of fMRI responses averaged over all voxels in the V1 ROI defined in panel A. Dotted grey lines, trial onsets. fMRI data are not artifact-corrected in this figure. **C**. Fourier transform of time course in panel B. Open red circle, amplitude of the frequency response at the task frequency. Open black circles, the rest of the frequency response. The largest frequency component of the TRR was at the task frequency.

### Head movement, cardiac activity, and respiration did not give rise to task-related fMRI responses

The influences of head movement, cardiac activity and respiration on the fMRI signal were removed from the data using the following sequence: ME-ICA (which reduces head movement artifacts (13–15)), global signal regression (6, 12), and projecting each movement/physiological regressor out of the resulting fMRI data. The maximum amplitude frequency component in the artifact-corrected time course (Fig. 3A) remained 1/15 Hz (i.e., the task frequency), though the amplitude at this frequency was reduced slightly by the artifact correction. In the following sections, all analyses were performed on artifact-corrected time series.

**Fig. 3.**
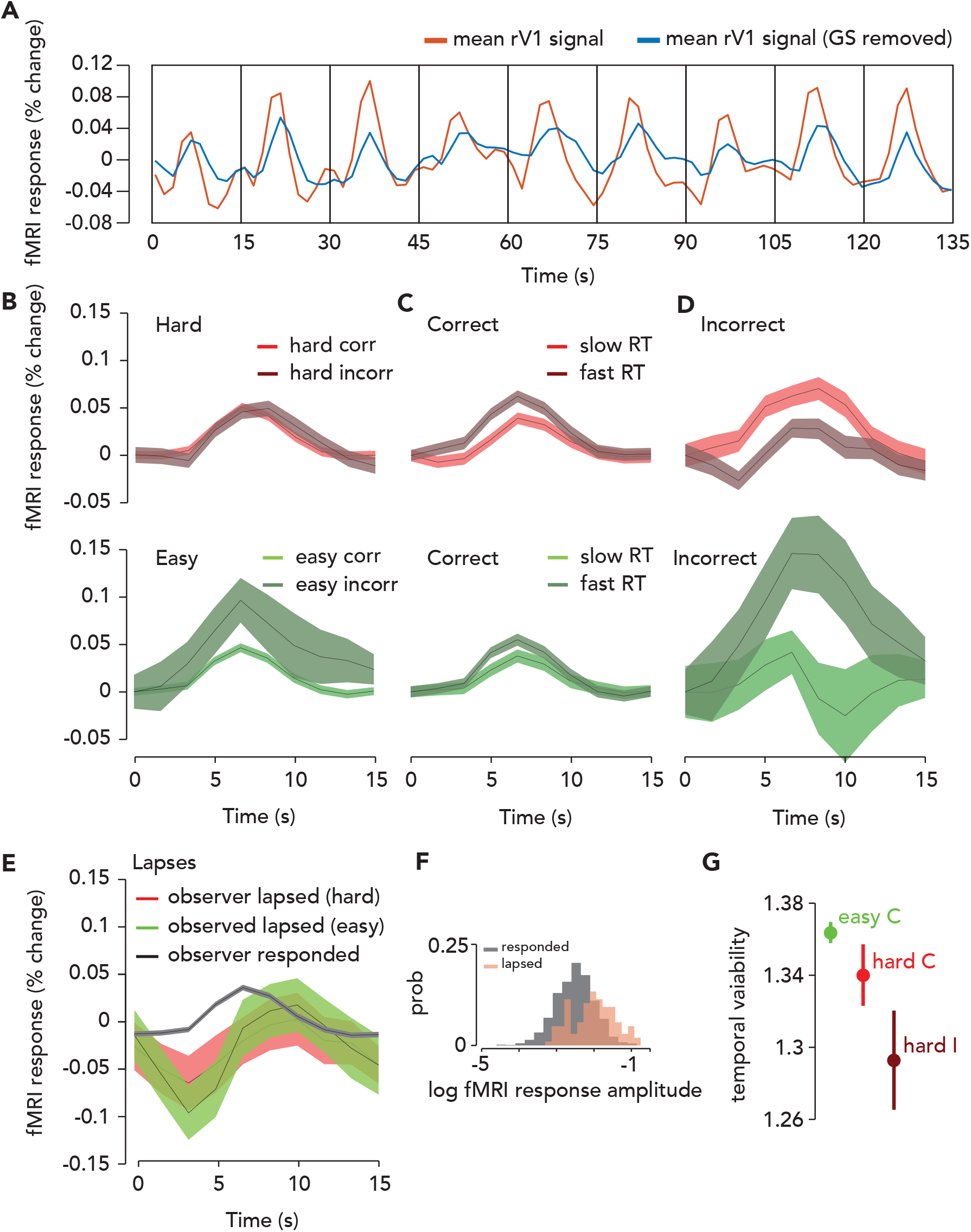
The amplitude and timing of the TRR were modulated by task difficulty and behavioral performance. **A**. To estimate the TRR, we projected the global signal (mean fMRI signal across all voxels in the brain) out of each voxel’s fMRI response in ipsilateral (right) V1 (red, average across V1 voxels before correction). We then averaged over all voxels in V1 (blue). **B**. TRR amplitude was modulated by task difficulty and behavioral accuracy. Average TRRs across trials and participants (N = 9) for hard and easy runs, correct vs. incorrect trials. Red colors, hard runs. Green colors, easy runs. Light colors, correct trials. Dark colors, incorrect trials. Error surface, two SEM across observers. **C**. TRR amplitude was modulated by reaction time on correct trials. Same format as panel B, but just for correct trials, and light colors indicate trials with slow reaction times (above median of the RT distribution), dark colors indicate fast reaction times (below median RT). **D**. TRR amplitude was modulated by reaction time on incorrect trials. Same format as panel C, but for incorrect trials. **E**. TRR amplitude was modulated by lapses (missed response trials). Black curve, average TRR across all trials on which the observer made a button press response, irrespective of type. Red, lapsed trials on hard runs. Green, lapsed trials on easy runs. **F**. Distribution of (log) fMRI amplitude (measured as std of the signal on each trial), across trials for lapse vs. response trials (orange vs. black). **G**. Temporal variability of the TRR is modulated by difficulty and behavioral accuracy. Circle, mean temporal variability (measured as the circular std of the Fourier phase at the task frequency of 1/15 Hz) across observers (N = 9). Error bar, two SEM. Green, easy correct trials. Light red, hard correct trials. Dark red, hard incorrect trials. Easy incorrect not shown for visualization purposes because it is far below the y-limit of the graph, and the error bar is large (easy incorrect temporal variability: mean, 0.52; SEM, 0.36).

### Task-related fMRI responses scaled with task difficulty

An observer needs to be alert to do well on a more challenging task. Therefore, we predicted that arousal would be higher on hard than easy trials, and that if TRRs reflect arousal, their amplitude would scale with task difficulty. Task-related responses in V1 were higher in amplitude on hard than easy trials (Fig. 3B). To quantify this effect, we ran a GLMM which included the following predictors: the convolution of each observer’s hemodynamic response function with impulses time-locked to (1) trial onset (which coincided with the stimulus presentation), (2) button press, and (3) a time-on-task boxcar in-between trial onset and button press; (4) task difficulty (easy or hard); (5) behavioral accuracy; (6) a prediction for the influence of cardiac activity (7) and respiration (8) on the fMRI signal (following refs (16, 17)); (9) the translational and rotational movement of the head, consisting of 6 parameters (see Methods). We also included terms that accounted for differences in the amplitude of the fMRI signal across observers, effectively normalizing the data across observers (11).

We focused our analysis on the interactions of task difficulty and accuracy with the linear combination of the three fMRI predictors (hereafter referred to as the “combined fMRI predictor”), which were individually labelled fMRI_TO, fMRI_BP, and fMRI_ToT, corresponding to trial onset, button press, and time-on-task locked fMRI activity, respectively. It was essential to include in the model both difficulty and accuracy, as well as their interactions with each other and with the three fMRI predictors to isolate the effect of task difficulty from behavioral performance, which is also linked to arousal (10, 18, 19). The rationale is that we wanted to be as liberal as possible in our assumptions about which task events the TRR is responsive to, and also following similar models commonly used to link arousal to pupil size (20, 21) and task-related neural activity to the fMRI signal (22) (see Discussion for more details). The full statistical results, with p-values for every predictor and interaction, are reported in the Supporting Information (File S1). The interaction of task difficulty with the combined fMRI predictor was significant (p = 6.44e-7, F = 10.53, N = 16,399; (fMRI_TO+fMRI_ToT+fMRI_BP):difficulty), demonstrating that the amplitude of the TRR was larger when the task was more challenging.

### Task-related fMRI responses scaled with behavioral accuracy on individual trials

Task-related responses were higher in amplitude for trials on which observers made an incorrect behavioral report (Fig. 3B). We hypothesized that the auditory feedback, which immediately followed the observer’s button press and indicated whether the response was correct or incorrect, may have surprised observers on the infrequent incorrect trials, and hence increased arousal and thereby modulated the amplitude of the TRR (10). Indeed, the interaction of behavioral accuracy with the combined fMRI predictor was significant in the GLMM (p = 0.005; F = 4.24; N = 16,399; (fMRI_TO+fMRI_ToT+fMRI_BP):accuracy), as was their interaction with difficulty (p = 1.33e-4; F = 6.84; N = 16,399; (fMRI_TO+fMRI_ToT+fMRI_BP):accuracy:difficulty). The amplitude of the TRR was highest on easy incorrect trials, when we expected that observers would be most surprised by their own (infrequent) errors (10).

### Task-related fMRI responses scaled with reaction time and this relation was captured by linear summation of three task-related components

Arousal influences behavioral performance, including reaction time (10, 23), so we hypothesized that the amplitude of the TRR would scale with reaction time. For correct trials, regardless of task difficulty, TRRs were higher in amplitude when the reaction time was fast (Fig. 3C). For incorrect easy trials, this effect persisted and was even larger (Fig. 3D), whereas for incorrect hard trials, the effect reversed and TRRs were higher in amplitude when reaction time was slow.

Reaction time was represented in our model in two unique ways: first, by the latency between the trial onset and button press inputs, and second, by the duration of the time-on-task input (Fig. S1). We hypothesized that amplitude modulation of the TRR with reaction time (on individual trials) was determined by the sum of these three hypothesized inputs. If this hypothesis is correct, the model and data should exhibit a similar pattern of amplitude modulation with reaction time on average across trials and participants. To test this, we performed a simulation in which we (1) predicted fMRI activity in response to the trial onset, button press, and time-on-task inputs using the reaction times measured in the task, (2) averaged the simulated responses over all high and low RT trials (median split) and participants, crossed by difficulty and accuracy, and (3) compared the pattern of amplitudes between data and simulation. We found that some aspects of the data were captured best by a sole time-on-task input (“fMRI_ToT”) and other aspects were captured better by just the linear sum of the trial onset and button press inputs (“fMRI_TO+fMRI_BP”), and overall the data lay somewhere in-between these two extremes (Fig. S2; see Discussion for motivation for focusing on these two cases). Specifically, the fMRI_TO+fMRI_BP simulation captured the pattern of amplitude modulation with reaction time on correct trials as well as the overall amplitude modulation with accuracy, and the fMRI_ToT simulation better captured RT-dependent amplitude modulation seen on incorrect trials. This validates our use of a model with all three inputs, which can interpolate between these two extremes in a flexible way. These results demonstrate that amplitude modulation of the TRR with reaction time may arise from linear summation of three task-related components, a model commonly used for linking arousal to pupil size (19–21, 24).

### Task-related fMRI responses were large and delayed on lapsed trials

A previous study found that TRRs in macaque V1 were largest when the animal disengaged from the task (3), suggesting that TRRs may become largest, paradoxically, at very low arousal levels. We estimated TRRs in human V1 on lapse trials, defined as trials on which the observer didn’t respond. We found that TRRs were much larger in amplitude on lapse than response trials, and this effect was stronger on easy than hard trials (Fig. 3E,F). TRRs were also delayed on lapse trials, peaking at 10 s after trial onset, rather than the typical 6 s on response trials. The TRR also displayed a more prominent decrease on lapse trials (Fig. 3E,F).

### Task difficulty modulated trial-to-trial temporal variability in task-related fMRI responses

Reward magnitude is known to modulate the timing precision, as well as the amplitude of TRRs (3, 6). We tested whether task difficulty similarly modulates timing precision. We measured temporal variability (across trials) in the TRR as the circular standard deviation of the Fourier phase as the task-onset frequency (1/15 Hz). Timing precision was modulated by both task difficulty and behavioral accuracy (Fig. 3G), being highest for easy incorrect trials, next highest for hard incorrect, followed by hard correct, and lowest for easy correct (compare with Fig 5. In ref (6)). These results indicate that the modulation of temporal variability of the TRR is not specific to changes in reward (3, 6), but rather may be observed in a range of task contexts in which arousal levels vary.

### Pupil size, respiration, and cardiac activity were similarly modulated by task difficulty and behavioral performance

If modulation of TRRs with task difficulty and behavioral performance reflect changes in arousal, we should observe that common physiological measures of arousal are also similarly modulated. The fMRI time series that we analyzed was corrected for artifacts caused by head movements, cardiac activity, and respiratory activity. Hence, the following analysis instead tests the idea that a common arousal process modulates both the TRR and physiological processes, such as cardiac and respiratory activity, independent on the artifactual impact they have on the BOLD signal. We had a group of observers (N = 5) perform the same behavioral task (with shorter 4 s trials) outside the fMRI scanner, while pupil size was measured with an infrared eye-tracker. The amplitude of the task-evoked pupil response was modulated by accuracy and task difficulty (Fig. 4A,B; permutation test, p < 0.05 for both comparisons, N =5). The observed modulation (incorrect > correct, with easy showing a bigger effect than hard) was similar to the fMRI results.

**Fig. 4.**
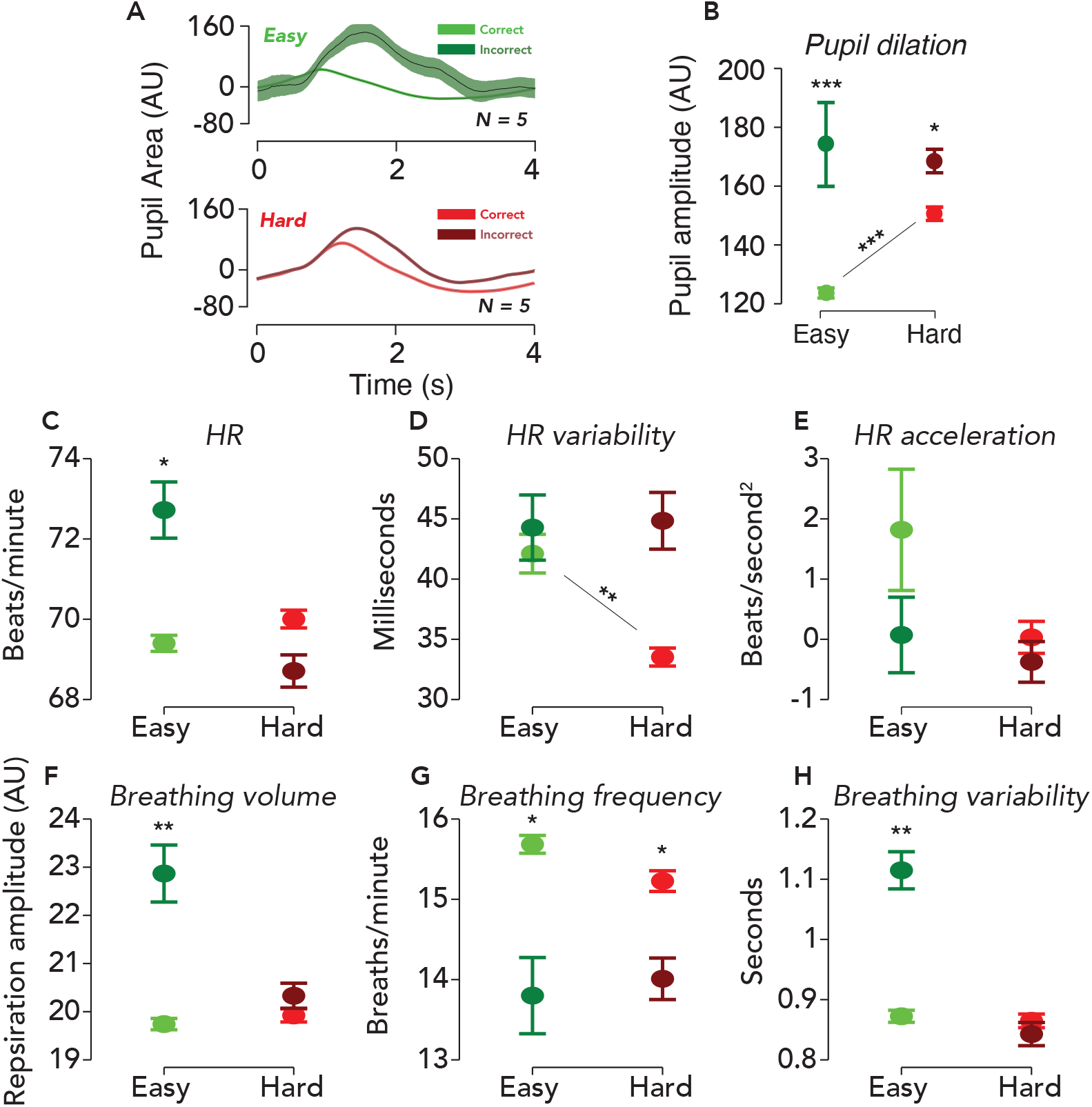
Physiological responses were similarly modulated by difficulty and behavioral accuracy. **A**. Task-evoked pupil responses. Solid black lines, average task-evoked pupil response across trials and observers. Error surfaces, two SEM, across observers (N = 5). Green, easy runs of trials. Red, hard runs. Light colors, trials with a correct behavioral report. Dark colors, incorrect. All aspects of the pupil and fMRI protocols were identical, except the ISI was 3.8 seconds in the former and 14.8 seconds in the latter. **B**. Amplitude of pupil dilation, measured as the standard deviation of the task-evoked pupil response time course. Error bar, two SEM. Asterisks, *: p < .05, **: p < .01, ***: p < .001 (permutation tests; one-tailed). Test statistic, the difference in the mean response amplitude for easy incorrect vs. correct trials, hard incorrect vs. correct trials, or hard correct vs easy correct trials. **C-H**. Physiological measurements collected during fMRI scanning were modulated by task difficulty and behavioral accuracy. Circle, average of physiological measure across time, trials, and participants. Error bar, two SEM (N = 9). Same format as panel B, with asterisks indicating result of permutation test.

Heart rate, heart rate variability, and acceleration, as well as respiration volume, frequency, and variability (see Methods for definitions), all of which are known to be influenced by arousal (i.e., via sympathetic and parasympathetic nervous system activity; see Methods), also exhibited similar modulations (Fig. 4C-H). We attempted a GLMM approach to predict the physiological measures from task and behavioral variables, however standard checks of the residuals and fitted values indicated violations of the model’s linear-Gaussian assumptions. Thus, we instead used non-parametric permutation tests to analyze dependency of the physiological measures on task difficulty and accuracy (see Methods). We found that for easy runs of trials, all physiological measures except heart rate variability and heart rate acceleration were significantly modulated by accuracy (easy incorrect > easy correct; heart rate, p = 0.032; heart rate variability, p = 0.719; heart rate acceleration, p = 0.507; respiration volume, p = 0.004; respiration frequency, p = 0.035; respiration variability, p = 0.005). For hard runs of trials, only breathing frequency was significantly modulated by accuracy (hard incorrect < hard correct; p = 0.015; comparisons for all other physiological measures: hard incorrect > hard correct; p > 0.05). Only heart rate variability was significantly modulated by difficulty, analyzing correct trials only (hard correct < easy correct; p = 0.005; comparisons for all other physiological measures: hard correct > easy correct; p > 0.05). Physiological measures of arousal were modulated by task difficulty and behavioral performance in a similar manner to the TRR, suggesting arousal as a common driver.

### Task-related fMRI responses were progressively weaker throughout the visual cortical hierarchy

We measured TRRs in areas V1, V2, and V3, always ipsilateral to the visual stimulus. TRRs were highest in amplitude in V1, and became progressively weaker in V2 and V3 (Fig. 5A-C). Amplitude modulation with difficulty and accuracy was found in each region, with the strongest modulations in V1, and becoming smaller in amplitude at progressively higher levels of the visual cortical hierarchy. TRR amplitude was measured non-parametrically as the standard deviation of the response time course on a single trial and summarized visually in Fig. 5B. The temporal variability of the TRR, on the other hand, was similar between V1, V2, and V3 (Fig. 5D);

**Fig 5.**
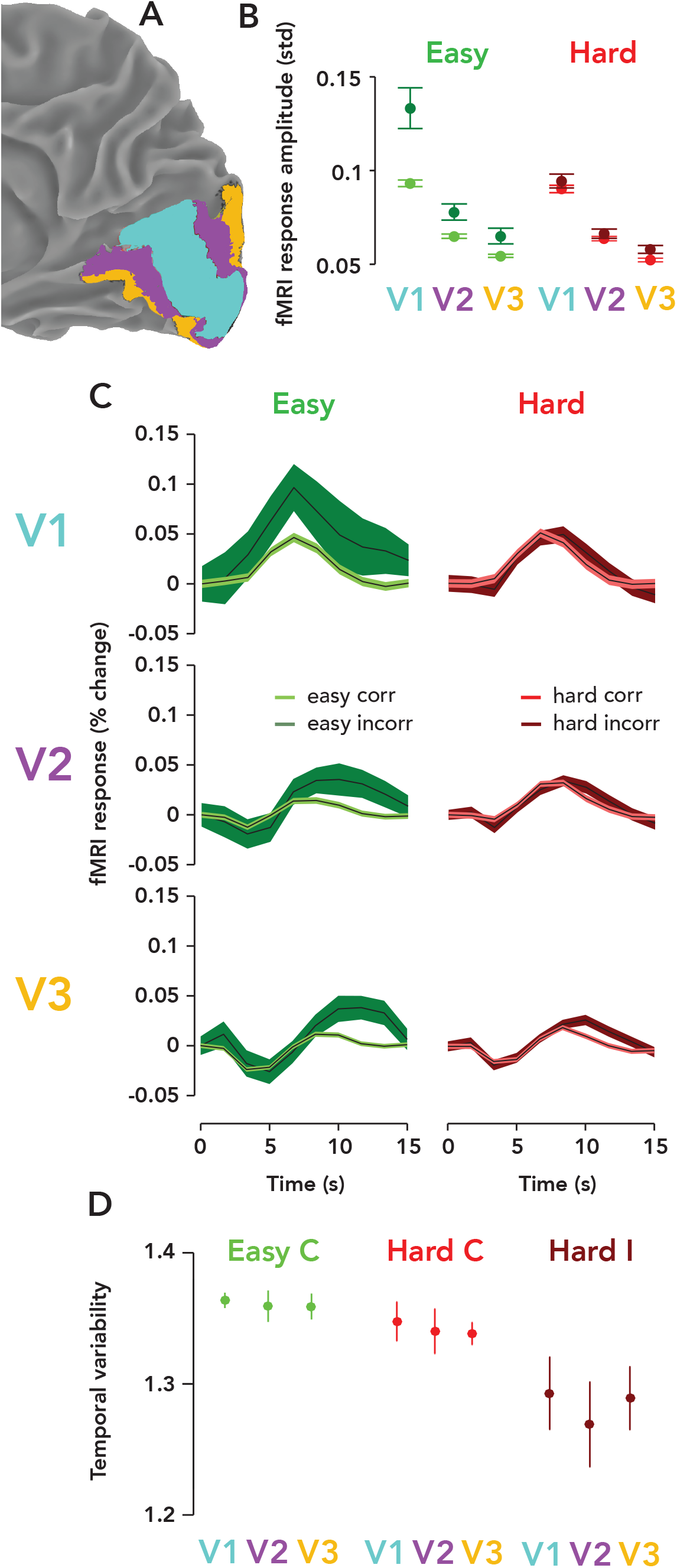
Task-related responses were progressively weaker ascending the visual cortical hierarchy. **A**. Posterior third of the right hemisphere (ipsilateral to stimulus), Observer 8, with atlas-defined V1, V2, and V3 highlighted in three different colors (cyan, purple, orange, respectively). **B**. fMRI response amplitude, measured as the standard deviation of the time course on each trial, scaled with task difficulty and behavioral accuracy in V1-V3. Weaker responses and modulations were observed in V3 than V2, and in V2 than V1. Circle, mean fMRI response amplitude across observers (N = 9). Error bar, two SEM. Green, easy runs. Red, hard runs. Light colors, correct trials. Dark colors, incorrect trials. **C**. Mean TRRs across observers for different trial types and ROIs. Same format as Fig. 3B. **D**. Temporal variability is similar across V1, V2, and V3. Same format as Fig. 3G.

## Discussion

We observed large fMRI responses in human early visual cortex that were spatially widespread, independent of visual stimulation, and tightly entrained to trial timing. These TRRs, measured in V1 in the hemisphere ipsilateral to the stimulus, were larger in amplitude when the task was more difficult, and on trials on which observers made an erroneous response, failed to respond, or were faster to respond. Neither the TRR nor its modulation with task difficulty or behavioral performance was explained by common fMRI-BOLD artifacts, i.e., changes in head movement, cardiac activity, and respiration, nor by the fMRI “global signal.” The timing precision of the TRR was also modulated by task difficulty and behavioral performance, complementing similar findings from previous studies that manipulated reward (3, 6). Contrary to the consensus of opinion that the fMRI-BOLD signal in V1 primarily reflects stimulus-evoked and attentional neuronal activity, we found evidence that the BOLD signal also contains a component that reflects the demands of a task and behavioral performance on the timescale of individual trials.

TRRs in early visual cortex are modulated by task difficulty, reward, and behavioral performance. Two previous studies (3, 6) examined the effect of reward magnitude on TRRs in early visual cortex, and found that larger rewards led to increased timing precision and increased response amplitude on individual trials, as well as larger trial-averaged response amplitude. Roth et al (6) found that multiple measures of trial-to-trial variability in TRRs (i.e., timepoint, amplitude, and temporal variability) changed with reward. The authors used computational simulations to show that simultaneous increases in trial-averaged TRR amplitude with reward and decreases in the three measures of variability were consistent with only one possibility: changes in trial-to-trial temporal jitter. That is, averaging over many temporally jittered responses led to a lower amplitude trial-averaged response whereas averaging over many temporally aligned responses led to the opposite. And likewise, more temporal jitter also generated increases in all measures of trial-to-trial variability (timepoint, amplitude, and temporal variability). However, these results don’t preclude additional changes in trial-to-trial response amplitude with reward, but instead suggest that a portion of the trial-averaged response amplitude cannot be predicted by trial-to-trial amplitude alone. Indeed, Cardoso et al. found that both the trial-to-trial amplitude and timing of TRRs in macaque V1 were modulated by reward (3). Our results provide a complementary account. We found that trial-to-trial amplitude, trial-averaged amplitude, and temporal variability of the TRR were all modulated by task difficulty and behavioral performance. Temporal precision was greater for unexpected task events (i.e., incorrect feedback tone on an easy run of trials), consistent with previous findings showing that temporal precision scaled with arousal (3, 6). Taken together, the evidence suggests that reward, task difficulty, and behavioral performance modulate both the timing precision and amplitude of TRRs.

We found that TRR amplitude was modulated by reaction time on a trial-to-trial basis. It is well known that arousal influences reaction times, with a classic inverted-U relation (25), an effect that shows up in standard physiological measures of arousal, including pupil size (10) and cardiac activity (26). So if TRR amplitude reflects arousal, as we hypothesized, we would expect that it would be higher when an observer responds faster, which is what we found (on correct trials). Note that we assume that our task primarily operates in the left half of the inverted-U, because even the hard task is rather boring, so arousal levels are never very high (10). Previous fMRI studies offer an alternative, but complementary perspective on the relationship between reaction time and TRR amplitude. A previous study measured the TRR in human V1 in a visual detection task (5) with either an immediate or delayed behavioral report (delay, ~9 seconds). In the delayed condition, there were response peaks following both trial onset and the behavioral report. In peripheral V1, far from the representation of the stimulus, the amplitude of fMRI activity was lower for a delayed than immediate report. This indicates that for an immediate report, there were also two components of the TRR — one for the trial onset and one for the response — but because they were so close in time, their impulse responses summed to a unimodal response. We might assume that two such ‘blurred-together’ responses were also present in our experiment, in which observers responded immediately after the stimulus presentation. It would follow then, that differences in reaction time should affect this summation and thereby modulate the amplitude of the unimodal TRR — slower RTs associated with lower fMRI amplitudes (Fig. S1A, Fig. S2B). Conversely, it could be that the amplitude of the TRR reflects time-on-task, i.e., a sustained input between trial onset and reaction time, often modeled as a boxcar (22). If true, TRR amplitude would be higher for trials with slower RTs (Fig. S1B, Fig. S2C). And a third possibility is that all three inputs are present, at trial onset, at the response time, and time-on-task. This scenario would produce some effect in-between these two extremes, which depends flexibly on the relative strengths of the three inputs. A model similar to this, with three inputs, is commonly used to predict the task-evoked pupil response from task structure (10, 20, 21). To be liberal in our assumptions, we used a GLMM with these three putative inputs — fMRI_TO, fMRI_BP, and fMRI_ToT — as well as their combinations. We found that the combination of all three inputs best explained the results (i.e., the interactions of the combined fMRI predictor with task and behavioral variables were significant in the model). This agrees with the results of many studies linking arousal and pupil size, which used a similar model (19–21, 24). Differences in reaction time are associated with changes in the timing and variability of phasic locus coeruleus (LC) activity (27). Thus, our results suggest that the influence of reaction time on TRR amplitude is mediated by a LC-NE arousal process.

The origin and control of the TRR in early visual cortex remain unclear. One possibility is that they are driven by an intrinsic vascular mechanism rather than a local neuronal one (1–3, 28–31). This putative mechanism, vasomotion or oscillation in tone of blood vessels independent of cardiac and respiratory activity, occurs in cerebral arteries and is influenced by arousal (32–34). More broadly, cerebral blood flow is complex and modulated by arousal via multiple cellular mechanisms (28, 35), including activity of astrocytes (36–39), pericytes (40, 41), endothelial cells (42), and smooth muscle cells (43). Our fMRI data cannot reveal the origins of the TRR, but may provide some clues. We found that the amplitude of the trial-averaged TRR was highest in V1, and progressively smaller in V2 and V3 (Fig. 5A), whereas timing precision was similar among the three visual areas (Fig. 5D). On the other hand, both timing precision and response amplitude were modulated by task difficulty and behavioral performance within each visual area (Fig. 5A,D). The larger TRRs in V1 (versus V2 and V3) may arise from some non-neural factor, possibly the unique vasculature of V1, where there are two input arteries, the calcarine artery (branch of the posterior cerebral artery) and posterior temporal artery (branch of the middle cerebral artery). This would support the view that the TRR is purely vascular in origin (31) and may be modulated (via vascular mechanisms) by neuromodulatory activity arising in the brainstem. Another possibility is that the TRR is driven by subthreshold neuronal activity (18), which previous measurements (i.e., LFP, spiking) were insensitive to (1, 31). Yet another possibility is that the TRR is driven by activity in a relatively small subpopulation of neurons, not evident in LFPs or mean firing rates.

Our findings may be related to previous studies identifying a component of the fMRI-BOLD signal related to arousal, but further research is necessary to test if this component and the TRR are the same (44–47). One of these studies (47) found that the most responsive voxels measured were in the primary sensory cortices, including early visual cortex, and that these voxels alone sufficed for prediction of behavioral performance and an EEG measure of arousal. If future studies find that this fMRI component is equivalent to the TRR, it could lead to a productive unification of these two areas of research.

The amplitude of the TRR in V1 (mean across all voxels and trials, 0.32% change; median, 0.28% change; max, 6.20% change) is similar to that of stimulus-evoked and attentional hemodynamic responses (2, 6, 31, 48, 49). This raises concerns about fMRI task designs in which attention or other psychological variables are operationalized according to behavioral performance and/or task parameters. In such cases, TRRs, which are modulated by critical task variables and behavior, may mix with other fMRI signal components, and thereby confound analysis. Our results demonstrate that common pre-processing approaches (i.e., global signal regression, physiological ‘noise’ correction) do not remove the TRR from fMRI data. Furthermore, the TRR contains information about brain state that would otherwise typically be measured independently with pupil size and/or cardiac activity. Therefore, we suggest that the TRR should be separately modeled and removed from fMRI data in preprocessing, and then possibly analyzed as an independent measure of arousal. One potential approach for accomplishing this would be to run a 5-10 minute version of a task in which the visual stimulus is confined to one visual field, so that the TRR can be easily isolated and measured later (see ref (50) for a simple protocol for removing the TRR).

## Conclusion

Approaches to analyzing and interpreting fMRI data are largely based on studies of early visual cortex, where neural activity is time-locked to visual stimulation and the fMRI signal is well approximated by a linear transformation of that stimulus-evoked neural activity (4, 51). However, it has been known for over a decade that the fMRI signal in early visual cortex contains large components not predicted by local spiking or visual stimuli (1, 4, 5). One of these components, the task-related fMRI response, is widespread and entrained to task timing, and it covaries with physiological measures of arousal as well as reward magnitude (1–3, 6, 29, 30). In this study, we found that the TRR is also modulated across trials by task difficulty, accuracy, reaction time, and lapses. These findings demonstrate the existence of a component of the fMRI-BOLD signal in human early visual cortex that reflects arousal on the timescale of individual trials.

## Materials and Methods

### Observers

Experiments were conducted at two sites. fMRI, cardiac, and respiration measurements were acquired from thirteen observers at the Functional Magnetic Resonance Imaging Core Facility at NIH. Pupil data were acquired from five other observers (two males, three females) at NYU. All observers were healthy adults, with no history of neurological disorders and with normal or corrected-to-normal vision. For the fMRI experiment, four observers were missing either high-quality cardiac and/or respiratory data, so all of their data was excluded from all analyses. Experiments were conducted with the written consent of each observer. The consent and experimental protocol were in compliance with the safety guidelines for MRI research, and were approved by both the University Committee on Activities involving Human Subjects at New York University, and the Institutional Review Board at the National Institutes of Health.

### Experimental Protocol

Observers performed a two-alternative forced-choice (2-AFC) orientation discrimination task, reporting whether a stimulus was tilted clockwise or counter-clockwise relative to vertical (Fig. 1). The stimulus was a grating patch (spatial frequency, 4 cycles/°) multiplied by a circular envelope (diameter, 1.5°) with raised-cosine edges (contrast, 100%). The stimulus was equiluminant with the gray background, such that it would not evoke a luminance response in visual cortex or a light reflex response in the pupil. On each trial, the grating was flashed briefly in the lower right visual field (5° diagonal to fixation) and observers covertly attended to the stimulus. Stimulus location was determined by the recording chambers used in related monkey electrophysiological experiments (3), which are typically centered at ~4–5° eccentricity in the lower visual hemifield, along the diagonal, i.e., ~45° or 135° polar angle. The difficulty of the task was manipulated by changing the tilt angle of the grating. In blocks of easy trials, the grating was tilted ±20° away from vertical. In blocks of hard trials, the grating was tilted by a much smaller amount (typically ±1°), with an adaptive staircase (one-up, two-down) ensuring ~70–75% correct discrimination accuracy. Observers were instructed to fixate a central cross on the monitor throughout the experimental session. The color of the fixation cross indicated the difficulty of the current block of trials — red for hard and green for easy blocks. This ensured that the observer was always informed of the current task difficulty. Each block had a predictable trial structure with a fixed inter-stimulus interval. Each trial was 15 seconds long, starting with a 200 ms stimulus presentation and a 14.8 s inter-stimulus interval, during which the observer was instructed to make a key press response indicating the perceived orientation of the grating. Observers could respond at any time during the ISI. 99.6% percent of RTs were under 4 seconds and 86.2% were under 1 second (median RT: 552 ms). A long ISI was employed because it allowed us to better measure the full time course of the TRR. Tone feedback was given immediately following the behavioral response — a high tone indicated correct behavioral responses and a low tone indicated incorrect behavioral responses. There were 16 trials per run (240 sec), but the first trial was always removed in our analysis to allow the hemodynamic response to reach steady-state. Stimuli were generated using Matlab (MathWorks, MA) and MGL (52) on a Macintosh computer. Stimuli were displayed via an LCD screen. Observers viewed the display through an angled mirror (field of view: 70 × 39.5 degrees).

### fMRI Data Acquisition

Blood oxygenation was measured in occipital, parietal, and temporal cortex with fMRI (3T GE scanner, 32-ch coil, multi-echo pulse sequence with echo times of 14.2 ms, 30.1 ms, and 46 ms; 22 slices covering the posterior third of the brain; voxel size 3×3×3 mm; TR of 1.5 s). fMRI time series were motion corrected (53) and then the time series from the three echos were combined into a single time series using ME-ICA (14).

### fMRI Analysis

fMRI data were aligned with base anatomical scans from each observer using a robust image registration algorithm (53). The boundaries of visual areas V1, V2, and V3 were defined using an anatomical template of retinotopy (54), and then projected onto each individual volumetric data, so as to avoid any interpolation or smoothing of the fMRI time series data (55). To characterize the TRR in ipsilateral V1, we computed the amplitude and phase of the sinusoid at the task frequency (1/15 Hz), which best fit the fMRI time series. Fourier analysis was used as validation that the largest frequency component of the TRR was at the task frequency. For Fig. 5A, the fMRI response amplitude (and pupil response amplitude) was computed as the standard deviation of the response time course over a single trial.

### Cardiac, Respiratory, and Head Movement Signals

While in the scanner, cardiac activity was measured using a pulse oximeter (GE Medical Systems E8819EH), respiration was measured using a respiration belt (GE Medical Systems E8811ED), and head movements were tracked continuously using real-time head motion estimates implemented in AFNI (56). To obtain a precise estimate of the peak times in the cardiac signal, given the temporal undersampling of the pulse oximeter, we used linear extrapolation based on signal to the left and the right of each putative undersampled peak location, and solved for the missing peak as the intersection of the two best-fit lines. Heart rate was computed as the mean of the time series of inter-peak durations and converted into beats per minute. Heart rate variability was computed using the Root Mean Square of the Successive Differences (RMSSD) method (57), which acts as a high-pass filter, removing low frequency components of the time series of inter-peak durations, such as those related to respiration. Heart rate variability is a widely-used measure of parasympathetic nervous system activity (58), and is typically higher when a participant is more calm. Heart rate acceleration was computed as the mean of the first derivative of the time series of inter-peak durations, with units of beats/s2, so positive numbers reflect acceleration and negative numbers deceleration. Acceleration/deceleration of the heart beat is associated with changes in either parasympathetic (vagus nerve) or sympathetic nervous system activity (59). Likewise, there is both parasympathetic (vagus nerve) and sympathetic innervation of the lungs, and thus respiration can reflect arousal via autonomic nervous system activity (25). The respiration signal was mean-subtracted and converted into percent signal change. Its amplitude (volume) was obtained by taking the standard deviation of the time series on each trial separately. Respiration frequency was computed, on each trial, as the component with the maximum amplitude in the frequency response. People hold their breath more when more alert, leading us to predict that respiration frequency should be lower when arousal is higher (25). Respiration variability was computed in the same way as heart rate variability, without the peak extrapolation method, by finding the signal peak times and computing the RMSSD thereof. We included all of these physiological measures because they capture different components of arousal (e.g., reflecting sympathetic and/or parasympathetic nervous system activity), which may have different links with behavior and task demands. Head movement estimates were computed from the times series from the first TE of the multi-echo sequence, and then this estimate was used to motion correct the images from all three TEs (14). We used these estimates of the translational and rotational movement of the head in each dimension of movement (roll, pitch, yaw, y, x, & z) as predictors in the GLMM.

To analyze the influence of task difficulty and behavioral accuracy on physiological measures of arousal, we binned these physiological measures by trial type (difficulty x accuracy) and ran permutation tests to check for differences among trial types. Permutation tests were performed to assess the differences in the amplitude of task-evoked pupil responses as well. We computed a test statistic from the data, the difference in the mean physiological measure across trials between easy correct and easy incorrect trials, hard correct and hard incorrect trials, and easy correct and hard correct trials. For heart rate variability and breathing frequency only, the direction of the test was reversed, as we expected these measures to be lower when arousal is greater. We constructed a null distribution by concatenating the data from the two conditions, randomly shuffling the trial type labels (10,000 iterations), splitting the resulting array into two new groups (with the same sizes as the original two labeled groups), and computing the difference of the mean of these two new groups. For each test, the p-value of the test was equal to the proportion of samples in the null distribution greater than the test statistic. These “exact p-values” were then corrected for the finite number of permutations performed (60).

### Data Cleaning

The first trial of each run was removed. This was done for all data types: fMRI, physiological, head movement, and behavioral data. Trials on which the observer did not respond were removed and analyzed separately as “lapse trials”.

### General linear mixed model

Our aim was to test the hypothesis that task difficulty and behavioral performance modulate the amplitude of TRRs in V1. We used a standard general linear model (GLM) approach, with additional random intercepts and slopes for each observer to account for systematic differences in the fMRI signal across observers, i.e., a GLMM (11). We included all predictors of interest in the model: putative TRR components (trial onset [“fMRI_TO”], button press [“fMRI_BP”], and time-on-task [“fMRI_ToT”]) (22), task difficulty, behavioral performance, predictions of the fMRI signal evoked by respiration and cardiac activity, and head movement. The GLMM had 24 fixed effects coefficients, corresponding to the following predictors: the convolution of a parametric HRF (the sum of two gamma functions) (61) fit separately to each observer’s grand mean TRR with impulses at (1) trial onset and (2) button press, and with a (3) time-on-task boxcar between them, as well as (4) task difficulty, (5) behavioral accuracy, (6) predictions for fMRI activity evoked by respiration (16) and (7) cardiac activity (17), (8–13) the movement of the head in 6 dimensions (roll, pitch, yaw, y, x, and z) and (13–24) the interactions of all fMRI, task, and behavioral predictors (every combination). All predictors were downsampled (and appropriately low-pass filtered first, i.e., decimated) to the sampling rate of the fMRI signal. We also included one random intercept for each observer, and random slopes for the three fMRI predictors for each observer, for a total of 63 random effects coefficients. Each predictor was a vector, comprising a single value per trial, for all trials and all observers. All predictors were included in a single design matrix, which was used to predict the trial-to-trial TRR in ipsilateral (right) V1. Head movement, cardiac, and respiration predictors were included to partial out their influence on the fMRI signal.

We used the fitlme function in Matlab to fit the GLMM to the voxel-average TRR (rV1) time course. We tested the assumptions of the linear model by examining the residuals as a function of the fitted values, and found that there was no trend, just a large Gaussian-looking cloud centered at zero. The distribution of residuals appeared Gaussian and the qq-plot showed slight deviation from Gaussian behavior at the tails, but Gaussian behavior in the center of the distribution. The total R2 of the model was 5.6%, suggesting that the inclusion of the multiple interaction terms in the model didn’t cause overfitting. The total R2 of the model was low in comparison to linear models of stimulus-evoked fMRI activity, but nonetheless many of the regressors were statistically significant. Due to the paucity of studies that have modeled the TRR, it’s unclear what one might expect this R2 to be, given the intrinsic variability of the TRR and measurement noise. Next, we examined the full time course of the model prediction vs. the data, and observed that the random slopes we included for the fMRI predictors were necessary to capture sometimes large difference in the fMRI signal amplitude across observers. To compute p-values and F statistics for linear combinations of the predictors (e.g., the combination of the three fMRI inputs, and its interaction with difficulty and/or accuracy), we performed an F-test using the coefTest function in Matlab.

To build the respiration and cardiac fMRI predictors, we convolved the raw signal from the respiration belt with a canonical respiration response function, which effectively models the influence of respiration on the fMRI signal (16). We did the same with the cardiac (pulse oximeter) signal, based on ref (17).

For each observer, we fit six parameters of a parametric HRF (the sum of two gamma functions) (61) to their grand mean trial-averaged TRR using fminsearch in Matlab. We could only use the 16 seconds of the grand mean TRR that we had to fit the parameters of the parametric form, but we allowed the resulting parametric HRF to be 32 seconds long. We used this best-fit parametric HRF to build the three fMRI predictors (fMRI_TO, fMRI_BP, and fMRI_ToT) by convolving the HRF with the impulses or boxcar occurring every trial (22).

### Pupillometry

The experiment was repeated outside the scanner with an identical protocol, except with a shorter, four-second ISI (10). Pupil area was recorded continuously during the task using an Eyelink 1000 infrared eye tracker (SR Research Ltd., Ontario, Canada) with a sampling rate of 500 Hz. Nine-point calibration and validation were performed before each run. Blinks and saccades were removed from the pupil size time series and interpolated over (using piecewise cubic interpolation). Pupil size time series from each run were band-pass filtered with a 4th order zero-phase (i.e., “filtfilt” in Matlab) Butterworth filter with 0.03 Hz and 10 Hz cutoffs (10). Average pupil responses, time-locked to trial onset, were calculated for hard and easy runs, correct and incorrect trials separately. The SEM for each average time series was computed across all runs and observers.

## Acknowledgements

Thanks to Aniruddha Das and Jonathan Winawer for their comments on the manuscript. This research was supported by National Eye Institute grant R01-EY025330 (to D.J.H.), the National Eye Institute Visual Neuroscience Training Grant, T32-EY007136 (to C.S.B. through NYU), and the National Defense Science and Engineering Graduate Fellowship (to C.S.B.), and by the Intramural Research Program of the National Institutes of Health (ZIA-MH-002909) - National Institute of Mental Health Clinical Study Protocol 93-M-0170, NCT00001360.

## Author Contributions

Conceptualization: EPM, DJH

Formal analysis: CSB, SM

Funding acquisition: DJH, EPM

Investigation: CSB, MR, ZNR, SM, EPM

Methodology: CSB, ZNR, SM

Project Administration: EPM

Resources: EPM

Software: CSB, EPM

Supervision: DJH, EPM

Visualization: CSB

Writing – Original Draft: CSB

Writing – Review & Editing: CSB, MR, ZNR, SM, DJH, EPM

## Supporting Information

**Fig. S1.**
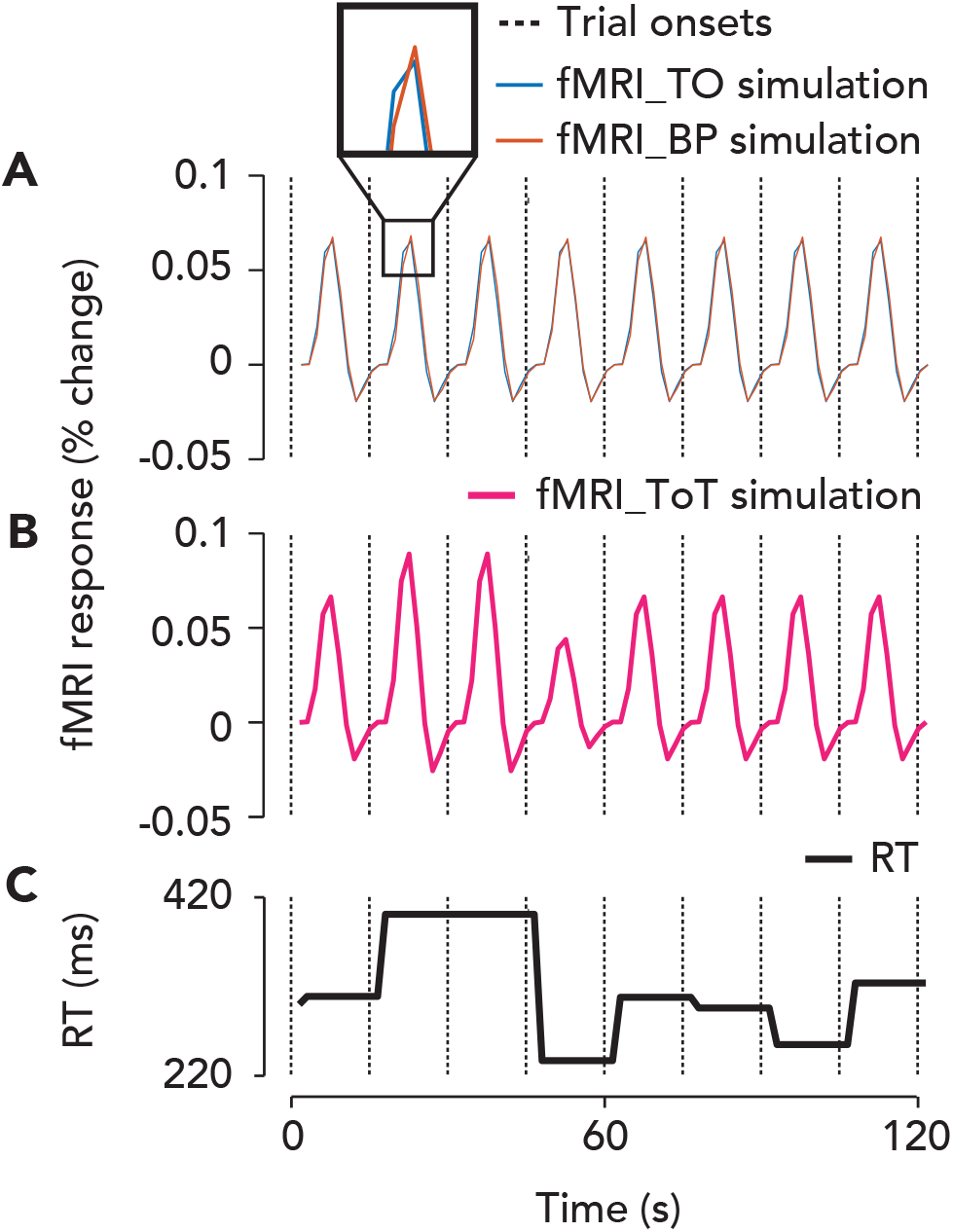
Trial onset, button press, and time-on-task fMRI predictors in the GLMM, and their dependence on reaction time. **A**. Blue curve, trial onset (“fMRI_TO”) locked fMRI response over eight trials. Red curve, button press (“fMRI_BP”) locked fMRI response. Dashed vertical lines, trial onsets. Inset, shows delay between the two responses, which varies from trial to trial depending on the reaction time. This delay will affect the amplitude of their sum, such that a longer delay will cause a lower amplitude summed response. **B**. Pink curve, time-on-task (“fMRI_ToT”) evoked fMRI response. Note that the time-on-task boxcar is longer in duration when the reaction time is slower, leading to a higher amplitude ToT-evoked fMRI response. **C**. Reaction time (RT) on each trial in milliseconds, taken from one observer’s data.

**Fig. S2.**
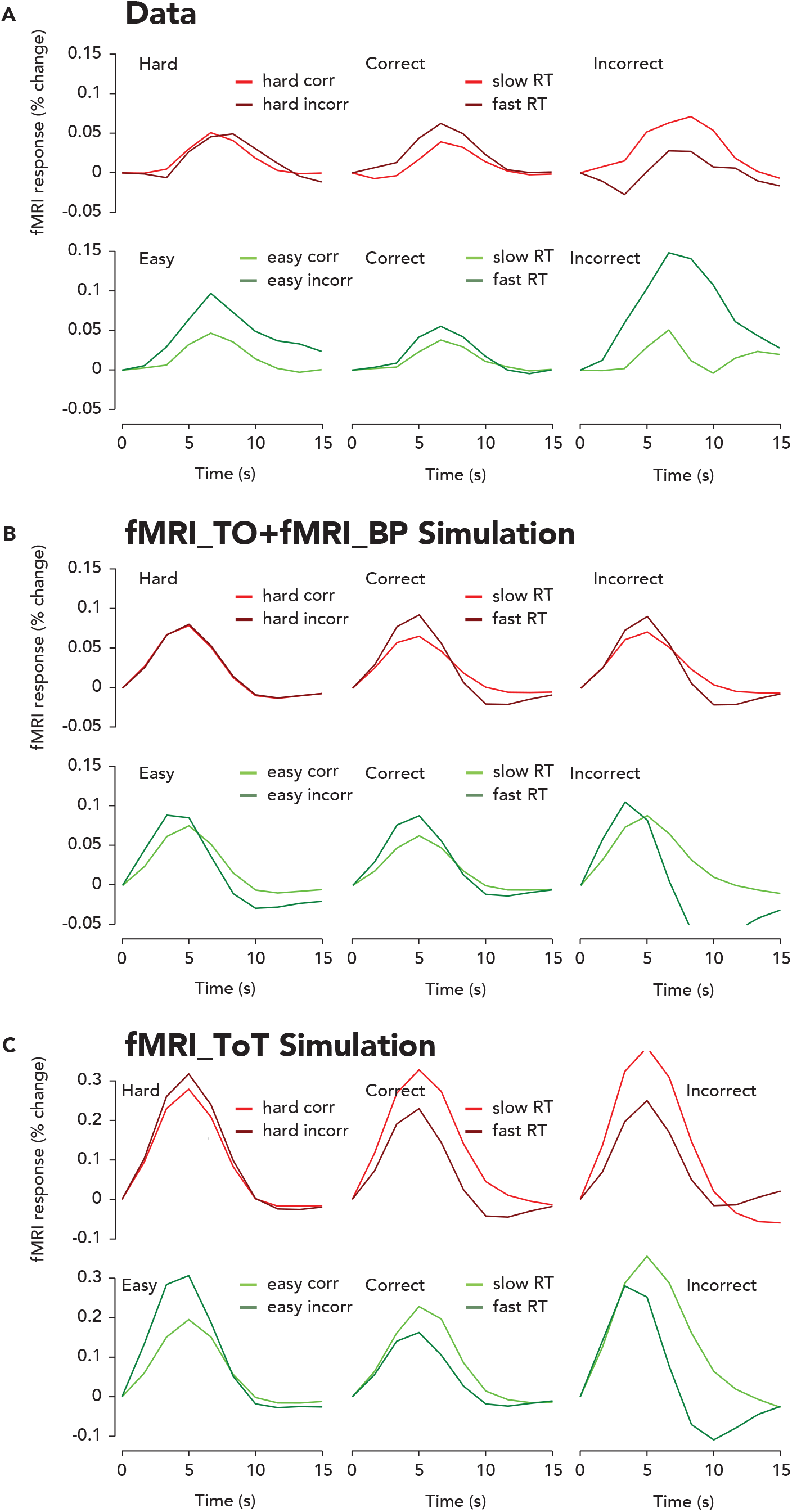
Amplitude modulation of the trial-averaged TRR with reaction time lies between two hypothetical extremes revealed by simulations. **A**. Trial-average TRRs for easy, hard, correct, incorrect, and fast and slow RT trials. Same format and color scheme as Fig. 3B-D, but without error bars for ease of comparison between data and simulation. **B**. Trial-average sum of the trial onset and button press locked fMRI responses, as plotted in Fig. S1A. Same format as panel A. Notice that the trial-averaged response is higher amplitude for fast than slow reaction times, consistent with shorter delays between the fMRI_TO and fMRI_BP inputs leading to a higher amplitude of the summed response. **C**. Trial-average time-on-task evoked fMRI response, as plotted in Fig. S1B. Notice that the trial-averaged response is higher amplitude for slow reaction times, consistent with a longer duration time-on-task input and greater temporal summation.

**File S1**. GLMM coefficients and p-values. See attached fileS1.txt file for full output from Matlab’s lme function.

